# Effect of organic zinc supplementation in hens on fertility from cryopreserved semen

**DOI:** 10.1101/2020.06.08.141390

**Authors:** Shanmugam Murugesan, Alagarsamy Kannan, Ramkrishna Mahapatra

## Abstract

Organic zinc supplementation in hen has been reported to improve fertility. The current study evaluated the effect of organic zinc supplementation in hens on fertility after insemination with cryopreserved semen. White Leghorn rooster semen was cryopreserved using 4% dimethylsulfoxide (DMSO) in 0.5ml French straws. Different semen parameters and fertility were assessed in post-thaw samples. White Leghorn hens were divided into 5 groups with 30 birds in each group. Each group was further divided into six replicates of five birds each. The control group was fed basal diet, other groups were fed with basal diet supplemented with 40, 60, 120 and 160 mg/kg organic zinc (zinc proteinate). After two weeks of feeding insemination was done in hens per vagina using thawed semen (200 million sperm/0.1 ml). Basal group hens were inseminated with fresh or cryopreserved semen and served as control groups. Sperm motility, live sperm, and acrosome intact sperm parameters were significantly (*p* < 0.05) lower in post-thaw semen samples. Fertility from cryopreserved semen was significantly (*p* < 0.05) lower and organic zinc supplemented hens had fertility similar to that of cryopreserved semen inseminated into basal diet group hens. In conclusion, organic zinc supplementation in hens does not improve fertility after insemination with 4% DMSO cryopreserved semen.

## 1. INTRODUCTION

Zinc is an important trace mineral in poultry that is involved in various biological and metabolic processes. Zinc is required for growth, normal functioning of reproductive and immune systems in chicken (Huang et al., 2019). Dietary supplementation of zinc in diet produced beneficial effects on laying performance, egg quality and antioxidant capacity in laying hens (Li et al., 2019). Zinc deficiency has been shown to affect hatchability in chicken (Blamberg et al. 1960). Zinc supplementation in chicken feed either had no effect on fertility (Stahl et al., 1986; Durmuş et al., 2004) or improved fertility and hatchability in addition to reduced embryonic mortality (Amen and Al-Daraji, 2011; Zhang et al., 2017; Li et al., 2019).

Fertility from cryopreserved semen is influenced by different factors such as breed/line of bird, cryoprotectant, cryopreservation protocol and presence of additives in the cryopreservation mixture (Donoghue and Wishart, 2000).

Studies have evaluated factors that improve fertility outcome from cryopreserved semen giving attention to male reproductive system. In chicken after insemination semen is stored in the sperm storage tubules (SST) up to three weeks and sperm are released from this storage site periodically so that the sperm move up the reproductive tract and fertilize the released ovum. The storage and release mechanisms of sperm are not fully deciphered (Sasanami et al., 2013). Turkey hens have been shown to influence the sperm penetration of inner perivitelline membrane and fertility and this effect is independent of sire (Christensen et al., 2006). Considering the foregone information it is not known whether manipulation of female reproductive system will improve fertility from cryopreserved semen. The aim of this study was to assess whether zinc supplementation in layer hens improve the fertility after insemination with cryopreserved semen.

## 2. MATERIALS AND METHODS

### 2.1 Experimental birds and husbandry

The experiment was carried out at the poultry farm of ICAR-Directorate of Poultry Research located in Hyderabad, India. The White Leghorn layers (IWH line) used in the experiment was housed in individual cages in an open-sided house. Feed and water were provided *ad libitum*. The experiment protocols were approved by the Institutional Animal Ethics Committee (IAEC/DPR/18/8).

### 2.2 Experiment

For the study, 150 White Leghorn layer hens of 38 weeks of age were selected and divided into 5 groups with 30 birds in each group. Each group was further divided into six replicates of five birds each. The five dietary treatments were Basal diet, Basal diet supplemented with 40, 80, 120 and 160 ppm organic zinc as zinc proteinate. The basal diet consisted primarily of corn and soybean meal (Table 1). Birds were subjected to 14 hrs of light per day. All hens were kept under the same managerial conditions. The supplemental trial period was for 10 weeks duration.

**TABLE 1.** Composition and nutrient levels of basal diet

| Ingredient | % |
| --- | --- |
| Maize | 61.46 |
| Soyabean meal | 24.74 |
| Deoiled rice bran | 0.42 |
| Stone grit | 10.90 |
| Di-calcium Phosphate | 1.50 |
| Salt | 0.50 |
| DL-Methionine | 0.10 |
| Trace minerals<sup>A</sup> | 0.10 |
| Vitamin premix<sup>B</sup> | 0.02 |
| Vitamin B Complex | 0.02 |
| Choline chloride | 0.10 |
| Toxin binder | 0.10 |
| Tylosine | 0.05 |
| Total | 100 |
| **Calculated values** |  |
| Metabolizable energy, MJ/kg | 11.33 |
| Crude Protein, % | 17.22 |
| Crude fibre, % | 3.32 |
| Lysine,% | 0.84 |
| Methionine, % | 0.37 |
| Methionine and cystine , % | 0.58 |
| Calcium, % | 4.56 |
| Nonphytate P, % | 0.47 |
| Zinc, mg/kg | 52 |

<sup>A</sup> Supplied (mg/kg diet):Mn, 24.2; Zn, 22; Cu, 5; Fe, 23.1; Se, 0.68; I, 1.9; Co, 0.21.
<sup>B</sup>Supplied (mg/kg diet): thiamin 1; pyridoxine, 2; cyanocobalamine, 0.01; niacin, 15; pantothenic acid, 10; a tocopherol, 10; riboflavin, 10; biotin, 0.08; menadione, 2; retinol acetate, 2.75; cholecalciferol, 0.06; choline, 650.

### 2.3 Semen collection and processing

Fifteen White Leghorn roosters aged 39 weeks were earlier trained to respond to abdominal massage technique (Burrows and Quinn, 1937) for collection of semen. Semen was collected randomly from the roosters on a day, pooled and kept on ice during the experiment. Collected semen was brought to the laboratory over ice in a covered thermocol box, evaluated and processed for cryopreservation. In the laboratory a portion of semen was diluted in a semen diluent (D (+)-glucose - 0.2 g, D (+)-trehalose dehydrate-3.8 g, L-glutamic acid, monosodium salt-1.2 g, Potassium acetate-0.3 g, Magnesium acetate tetrahydrate - 0.08 g, Potassium citrate monohydrate - 0.05 g, BES-0.4 g, Bis-Tris-0.4 g in 100 ml distilled water, pH 6.8; Sasaki et al., 2010) and was used for evaluation of semen quality parameters.

The pooled semen samples were initially evaluated for sperm concentration. The samples were diluted with cryoprotectant free diluent such that the sperm concentration was arrived at 4 million/μl. The samples were equilibrated at 5°C for 30 minutes and were diluted in 1:1 proportion with diluent containing 8% dimethyl sulfoxide (DMSO) so that the final concentration of DMSO was 4% and the final sperm concentration was 2 million/ μl in each treatment. The semen mixed with DMSO was immediately loaded into 0.5 ml French straws and sealed with polyvinyl alcohol powder. The filled straws were placed 4.5 cm above the liquid nitrogen (LN_2_) on a Styrofoam raft floating on LN_2_ in a thermocol box. The straws were exposed to nitrogen vapours for 30 minutes, plunged into LN_2_ and stored at −196°C until further use. Semen straws were stored for a minimum of seven days before evaluation. Cryopreserved semen after thawing at 5°C for 100 sec in ice water (Sasaki et al., 2010) was evaluated on nine different occasions for progressive sperm motility, live and abnormal sperm and intact sperm acrosome.

### 2.4 Sperm motility

Sperm motility was recorded as percentage of progressively motile sperm by placing a drop of diluted semen on a Makler chamber and examining under 20 x magnification. The percentage of sperm with normal, vigorous, and forward linear motion was subjectively assessed and scored.

### 2.5 Live and abnormal sperm

Percent live and abnormal sperm were estimated by differential staining technique using Eosin-Nigrosin stain (Campbell et al., 1953). Semen smear was prepared by mixing one drop of semen with two drops of Eosin-Nigrosin stain and air dried. Slides were evaluated under high power (100x) objective lens. All full and partially pink stained sperm were considered dead and unstained sperm as live. The percentage of live sperm was determined by counting at least 200 sperm. The same slides were used for estimating the abnormal sperm percent that were showing different morphological abnormalities.

### 2.6 Intact sperm acrosome

The intactness of sperm acrosome was assessed according to Pope et al. (1991). In brief, 10 μl of semen was mixed with 10 μl of stain (1% (wt/vol) rose Bengal, 1% (wt/vol) fast green FCF and 40% ethanol in citric acid (0.1 M) disodium phosphate (0.2 M) buffer (McIlvaine’s, pH 7.2-7.3) and kept for 70 sec. On a clean glass slide a smear of the mixture was made, dried and examined under high magnification (1000x). The acrosomal caps were stained blue in acrosome-intact sperm and no staining in the acrosome region of acrosome reacted sperm. A minimum of 200 sperm were counted in each smear sample and the percent acrosome intact sperm was calculated.

### 2.7 Fertility trial

Fertility trial was conducted using cryopreserved semen. After two weeks of feeding trial in hens insemination was carried out. In fresh semen insemination group and DMSO control groups 15 hens/treatment were inseminated whereas in all other zinc supplemented groups 20 hens/treatment were inseminated. Insemination was done twice at five days interval. The semen straws were thawed at 5°C for 100 sec in ice water (Sasaki et al., 2010) and inseminated into hen per vagina with sperm concentration of 200 million sperm/0.1 ml. For cryopreservation control the basal diet group hens were inseminated with cryopreserved semen. Freshly collected semen was inseminated into basal diet group hens with 100 million sperm/0.1 ml dose. Eggs were collected from second day of first insemination and stored at 15°C until incubation. The number of eggs incubated in different treatments ranged from 84 to 128. The eggs incubated at standard conditions in an automatic setter were candled on 18th day of incubation for embryonic development. Infertile eggs were broken open to confirm absence of embryonic development.

### 2.8 Statistical analysis

Data were analyzed using SPSS 16 software and *p* < 0.05 was considered significant. Percentage data were arcsine transformed and analyzed. Statistical analyses of semen parameters and fertility were done by one-way ANOVA with Tukey’s post hoc test.

## 3. RESULTS

The progressive sperm motility, live sperm and acrosome intact sperm parameters were significantly (*p* < 0.05) lower in post-thaw semen samples (Table 2). Fertility from cryopreserved semen was significantly (*p* < 0.05) lower in comparison to fresh semen insemination (Fig. 1). Fertility obtained in organic zinc supplemented hens was similar to that from cryopreserved semen inseminated in hens fed basal diet. No fertile eggs were obtained from 120 ppm organic zinc supplemented group.

**TABLE 2.** In vitro semen parameters in fresh and post-thaw White Leghorn semen cryopreserved using Sasaki diluent and 4% dimethyl sulfoxide.

| Parameters | Fresh semen | 4% DMSO |
| --- | --- | --- |
| Progressive sperm motility (%) | 65.56 ± 1.6<sup>a</sup> | 16.11 ± 1.62<sup>b</sup> |
| Live sperm (%) | 77.16 ± 2.8<sup>a</sup> | 21.28 ± 1.92<sup>b</sup> |
| Abnormal sperm (%) | 2.10 ± 0.17 | 2.19 ± 0.31 |
| Acrosome intact sperm (%) | 93.56 ± 1.4<sup>a</sup> | 87.91 ± 2.42<sup>b</sup> |

Values are mean±SE obtained from nine independent evaluations.
Figures bearing different superscripts in a row differ significantly ( $p < 0.05$ ).

**Figure 1.**
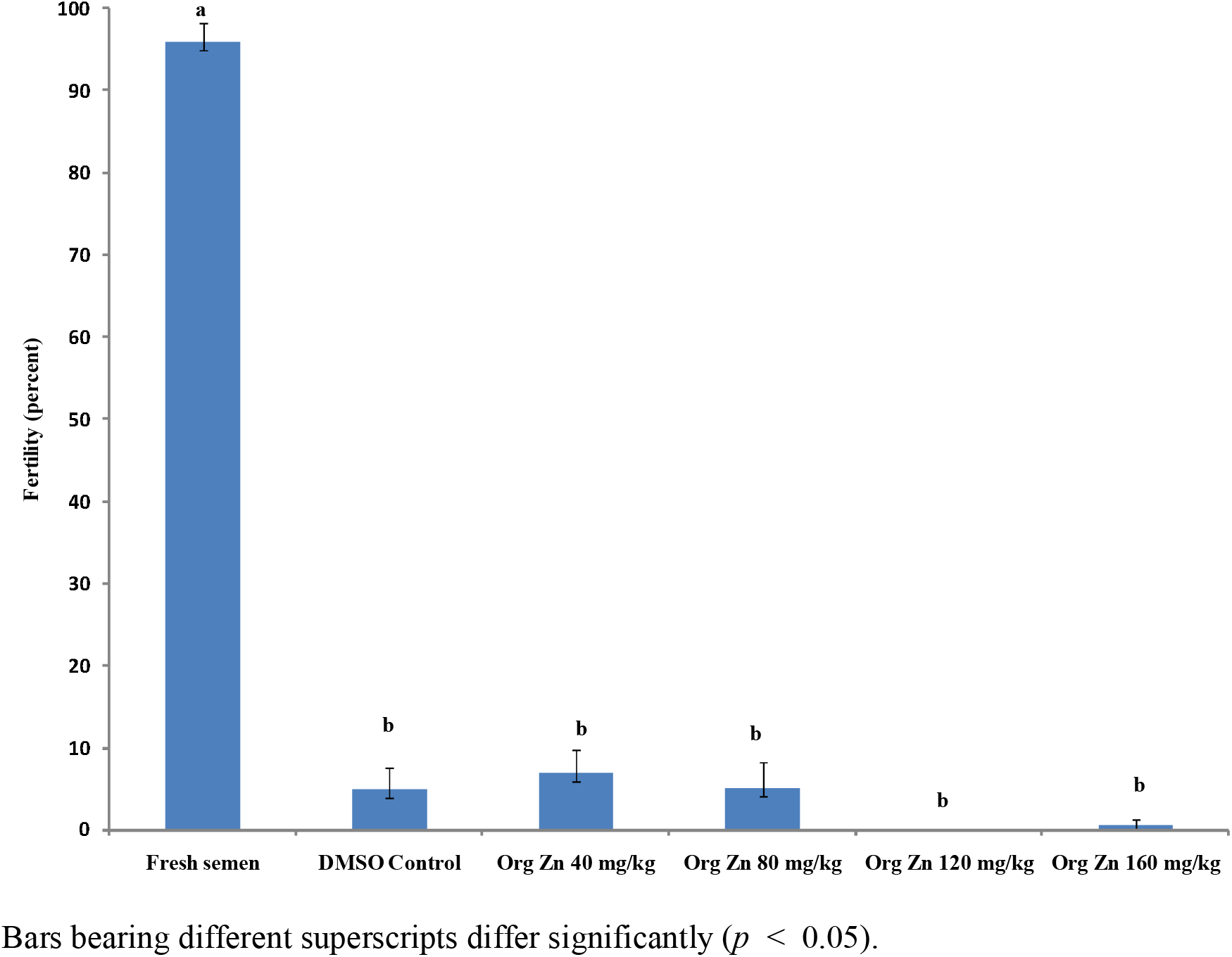
Fertility after insemination of cryopreserved semen in organic zinc supplemented White Leghorn hens

## 4. DISCUSSION

Fertility after insemination with cryopreserved semen in chicken is variable and research is undertaken to improve it by manipulating the cryopreservation protocols. The present study evaluated a novel way to improve fertility from cryopreserved semen through organic zinc supplementation in hens. The National Research Council recommends inclusion of zinc at 50 and 65 mg/kg diet for optimum productive and reproductive performance respectively (NRC 1994). A corn soybean diet should contain about 72 mg Zn/Kg of diet for obtaining good fertility and hatchability (Kidd et al., 1993). Hens fed corn-soy diet with 30 mg Zn/kg for 29 weeks resulted in decreased fertility and hatchability (Anshan, 1990). Thus a corn soybean diet should contain sufficient amount of zinc for optimum reproductive performance. The sperm egg penetration test and fertility was higher in zinc (100 mg/kg) supplemented broiler breeder hens (Amen and Al-Daraji, 2011). The reasons attributed for the higher results due to zinc supplementation were improvement in storage of sperm in SST and sperm motility. Zinc supplementation as zinc glycinate has been shown to improve fertility in comparison to basal diet and as well as with hens supplemented with zinc sulphate (Zhang et al., 2017). In addition to fertility zinc glycinate supplementation improved the antioxidant status in the birds. Li et al. (2019) had also reported higher fertility and hatchability in Chinese yellow feathered chicken supplemented with zinc (48-120 mg/kg). Cobb 500 broiler breeder hens supplemented with zinc oxide (60-120 mg/kg) had higher fertility at the later phase of laying period (Sharideh et al., 2016). These studies indicated that the supplemental zinc as well as source of zinc has an effect on fertility. However, few studies have reported no effect of dietary zinc supplementation on fertility. Zinc supplemented either as zinc oxide or zinc methionine did not improve fertility or hatchability in broiler breeders (Kidd et al. 1993). No effect of zinc supplementation either as zinc carbonate (40 mg/kg) or zinc sulphate (2000 mg/kg) on fertility was observed in White Leghorn layers (Stahl et al., 1986; 1990). Similarly no effect of zinc supplementation up to 210 mg on fertility was reported in brown egg layers (Durmuş et al., 2004). In the present study fertility from cryopreserved semen was not affected by organic zinc supplementation. This may be due to the source of zinc or the breed used in the study. Though zinc supplementation had no effect in the present study, future research should evaluate fertility from cryopreserved semen using minerals or compounds that has been shown to improve fertility in hens under normal conditions. Thus the unique reproductive physiology of female birds should be taken into consideration for improving the fertility outcome from cryopreserved semen in conservation programs of avian species.

In conclusion, organic zinc supplementation in the hen diet does not improve fertility from cryopreserved semen.

## AUTHOUR CONTRIBUTIONS

Experiment was designed, data were analysed, interpreted and manuscript prepared by all three authors. Feeding trials were conducted by A. Kannan and semen cryopreservation and fertility trials were conducted by M.Shanmugam.

